# A *cis*-encoded RNA interaction controls sub-operonic *spoVG* expression in *Staphylococcus aureus*

**DOI:** 10.64898/2026.09.18.752551

**Authors:** Raimy Lynch, Meja Pettersson, Pauline M.L. Coulon, Joanna M. Biazik, Jai J. Tree, Daniel G. Mediati

**Affiliations:** Australian Institute for Microbiology and Infection, University of Technology Sydney, Ultimo, NSW 2007, Australia; College of Science and Engineering, Flinders University, Bedford Park, SA 5042, Australia; Electron Microscopy Unit, University of New South Wales, Sydney, NSW 2052, Australia; School of Biotechnology and Biomolecular Sciences, University of New South Wales, Sydney, NSW 2052, Australia

## Abstract

Bacterial operons enable co-transcription of multiple genes, yet post-transcriptional mechanisms permitting differential regulation of individual genes within a polycistronic transcript remains incompletely understood. We demonstrate that a *cis*-encoded RNA-RNA interaction within the bicistronic *yabJ*-*spoVG* transcript forms an RNase III substrate that represses the transcription factor SpoVG in *Staphylococcus aureus*. Disruption of the *cis*-encoded regulation elevated monocistronic *spoVG* expression, reduced biofilm formation and cell wall peptidoglycan thickness, and increased sensitivity to cell wall-targeting antimicrobials in clinical vancomycin-intermediate *S. aureus*, consistent with phenotypes observed in an independent SpoVG overexpression strain. These findings uncover a previously undescribed mode of *cis*-regulatory control within a polycistronic operon and establish regulatory mRNA-mRNA interactions as a mechanism for sub-operonic gene regulation.

**SIGNIFICANCE STATEMENT:** Bacterial operons couple multiple genes into a single transcript, yet cells often adjust the output of individual genes. How this is achieved after transcription, and whether a transcript achieves this on its own has remained unclear. We show that the unusually long 5’ untranslated region of the *Staphylococcus aureus yabJ* mRNA base-pairs with the intercistronic region of the *yabJ*-*spoVG* transcript, repressing *spoVG* mRNA encoding a transcription factor. Disrupting the duplex increased *spoVG* and reshaped the SpoVG regulon. In a vancomycin-intermediate clinical isolate, this *cis*-regulation controls cell wall thickening, biofilm formation, and susceptibility to antimicrobials. *Cis*-encoded regulation within a polycistronic transcript provides a general route to gene-specific control.

## INTRODUCTION

*Staphylococcus aureus* is a remarkably adaptable multidrug-resistant opportunistic pathogen. Isolates of methicillin-resistant *S. aureus* (MRSA) frequently cause community-acquired and nosocomial infections, and account for up to 19% of bacteraemia cases [1, 2]. Patient treatment is now largely limited to last-line antibiotics, and the cell wall-targeting glycopeptides vancomycin and teicoplanin remain among the few options for MRSA infections [3]. However, treatment failure is increasingly common and is attributed to isolates with glycopeptide tolerance, refered to as vancomycin-intermediate *S. aureus* (VISA) (MIC = 4-8 μg/mL) [3]. VISA isolates commonly display thickened cell wall peptidoglycan, which likely limits uptake and diffusion of the antibiotic to the division septum at mid-cell [4, 5]. Multiple heterogeneous mechanisms leading to intermediate vancomycin resistance have been implicated, including post-transcriptional gene regulation [6–10] and varying loss-of-function mutations [3], demonstrating that the molecular basis of VISA remains incompletely defined.

Bacterial operons represent a fundamental strategy of gene regulation by coupling the expression of functionally-related genes in a polycistronic arrangement into a single transcript. This organisation is ubiquitous in bacterial transcriptomes with approx. half of all bacterial genes co-transcribed as part of an operon [11, 12]. Co-transcription does not, however, enforce uniform expression. For example, up to 43% of *E. coli* operons display differential expression of the individual cistrons, despite arising from a single transcript [11]. Such sub-operonic control can occur transcriptionally, through independent internal promoters and terminators [13], or post-transcriptionally, through differences in the stability of individual mRNA [14] and through *trans*-acting regulatory small (s)RNA that target individual cistrons or direct endonucleolytic cleavage within intercistronic regions [6, 15, 16]. By contrast, whether *cis*-encoded elements within a polycistronic transcript can themselves direct the differential expression of the co-transcribed genes remains unknown.

RNase III is the principal double-stranded RNA endoribonuclease in *S. aureus*, coordinating rRNA maturation and mRNA turnover. RNase III preferentially processes >22-nt RNA duplexes with cleavage favoured at GC/CG base-pair sites, and these duplexes can form in *cis* or in *trans* [17]. UV-crosslinked RNase III-RNA mapping has shown that RNase III is enriched within the 5’ and 3’ untranslated region (UTR) of mRNA transcripts [8, 18], reflecting a broader role for UTRs as post-transcriptional regulatory hubs governing transcript stability and translation [19]. In *S. aureus*, the *vigR* 3’UTR base-pairs in *trans* with the *dapE* mRNA to generate an RNase III processing site to repress *dapE* [20], while a separate RNA-RNA interaction with *isaA* (encoding a cell wall autolysin) is proposed to occlude an RNase III cleavage site to stabilise the *isaA* mRNA [8]. The combined *trans*-encoded regulation of the *vigR* 3’UTR mediates vancomycin tolerance in VISA. In a *cis*-acting counterpart, the 3’UTR of the *icaR* mRNA base-pairs with its own 5’UTR to create a substrate for RNase III processing [21], coupling transcript decay to post-transcriptional control within a single mRNA transcript.

In *S. aureus*, the *yabJ*-*spoVG* operon produces a bicistronic transcript under control of the alternative sigma B factor (σB) [22, 23]. YabJ, a distant homolog of RidA in *E. coli*, has no confirmed functional role in *S. aureus*, while *spoVG* encodes the principle effector of the operon and functions as a global transcription factor [24]. SpoVG preferentially binds DNA containing a TAATT_T/A_ sequence motif [25]. The regulon of SpoVG is large and multifaceted, controlling cell wall biogenesis [26], antimicrobial resistance [24], carbohydrate metabolism [27], and virulence pathways encompassing surface-associated proteins [28], exotoxin production [23], and capsular polysaccharide synthesis [29]. As a result, the physiological consequences of altering *spoVG* are wide-ranging. Deletion of *spoVG* (but not *yabJ*) reduced tolerance to glycopeptide and oxacillin treatment [24], and increased cell aggregation and biofilm development [30, 31]. In a murine subcutaneous skin infection model, the *spoVG* deletion produced significantly larger abscesses and higher bacterial burden [32]. In *Listeria monocytogenes*, deletion of *spoVG* produced a hypervirulent phenotype and increased resistance to the peptidoglycan-degrading antimicrobial lysozyme [33]. SpoVG is further repressed at the level of translation by the *trans*-encoded sRNA SprX, which occludes the *spoVG* ribosome-binding site without destabilising the bicistronic transcript [6]. Notably, the bicistronic *yabJ*-*spoVG* transcript is also processed to release monocistronic *yabJ* and *spoVG* mRNA through an unknown mechanism, suggesting that gene-specific control may be exerted from within the operon itself.

In this study, we demonstrate that an RNA-RNA interaction within the *yabJ-spoVG* transcript is the most highly abundant interaction associated with RNase III in our interactome of MRSA. We find that a *cis*-regulatory interaction occurs between the 5’UTR of *yabJ* and the intercistronic region of the *yabJ*-*spoVG* transcript, generating an RNase III substrate that represses both the biscistronic transcript and the monocistronic *spoVG* mRNA. Disrupting the interaction by single nucleotide polymorphisms (SNPs) significantly increased *spoVG* mRNA and remodelled the SpoVG regulon, altering genes involved in capsule synthesis, cell wall biogenesis, and pathogenesis. The interaction-disrupting SNP also reduced biofilm development and re-sensitised VISA to cell wall-targeting antimicrobials through a significantly thinner cell wall peptidoglycan. Inducible overexpression of SpoVG independently mirrored (phenocopied) the reduced cell wall peptidoglycan and reduced biofilm development observed, confirming that elevated SpoVG confers these phenotypes. Together, these findings uncover a *cis*-regulatory RNA-RNA interaction that enables post-transcriptional control of monocistronic *spoVG* within the bicistronic operon, highlighting a new layer of gene regulation in polycistronic systems with broad applications in manipulating bacterial gene expression.

## RESULTS

### The *yabJ*-*spoVG* transcript forms a mRNA-mRNA interaction

We previously captured the RNA interactome associated with the endoribonuclease RNase III (using RNA proximity-dependent ligation and sequencing, termed RNase III-CLASH) [8, 18] in the clinical MRSA ST239 isolate JKD6009 [34]. A total of 2,129 statistically significant RNA-RNA interactions were captured in replicate experiments conducted across two independent laboratories. By grouping the RNA duplexes into either sRNA-sRNA, sRNA-mRNA, or mRNA-mRNA interaction classes defined by the transcript boundaries of each hybrid read, we find that mRNA-mRNA interactions contributed the majority of the combined RNA interactome associated with RNase III (total 1,531 mRNA-mRNA interactions, **Figure 1A**), suggesting that mRNA can exert post-transcriptional regulatory functions. Among the RNA-RNA interactions profiled, we recovered 438 statistically significant hybrid counts within the bicistronic *yabJ*-*spoVG* transcript, contributing to the highest read count of mRNA-mRNA interactions recovered (*n*=8, **Figure 1A** and **Supplementary Figure 1A**) [8]. Plotting hybrid read count against the predicted free energy of hybridisation (ΔG) determined using RNAduplex for each RNA-RNA interaction placed *yabJ*-*spoVG* amongst the most stable (ΔG_hybridisation_ = −38.6 kcal/mol) and comparable to *bona fide* regulatory interactions, including regulatory mRNA-mRNA interactions previously characterised in *S. aureus* (**Figure 1B**).

**Figure 1.**
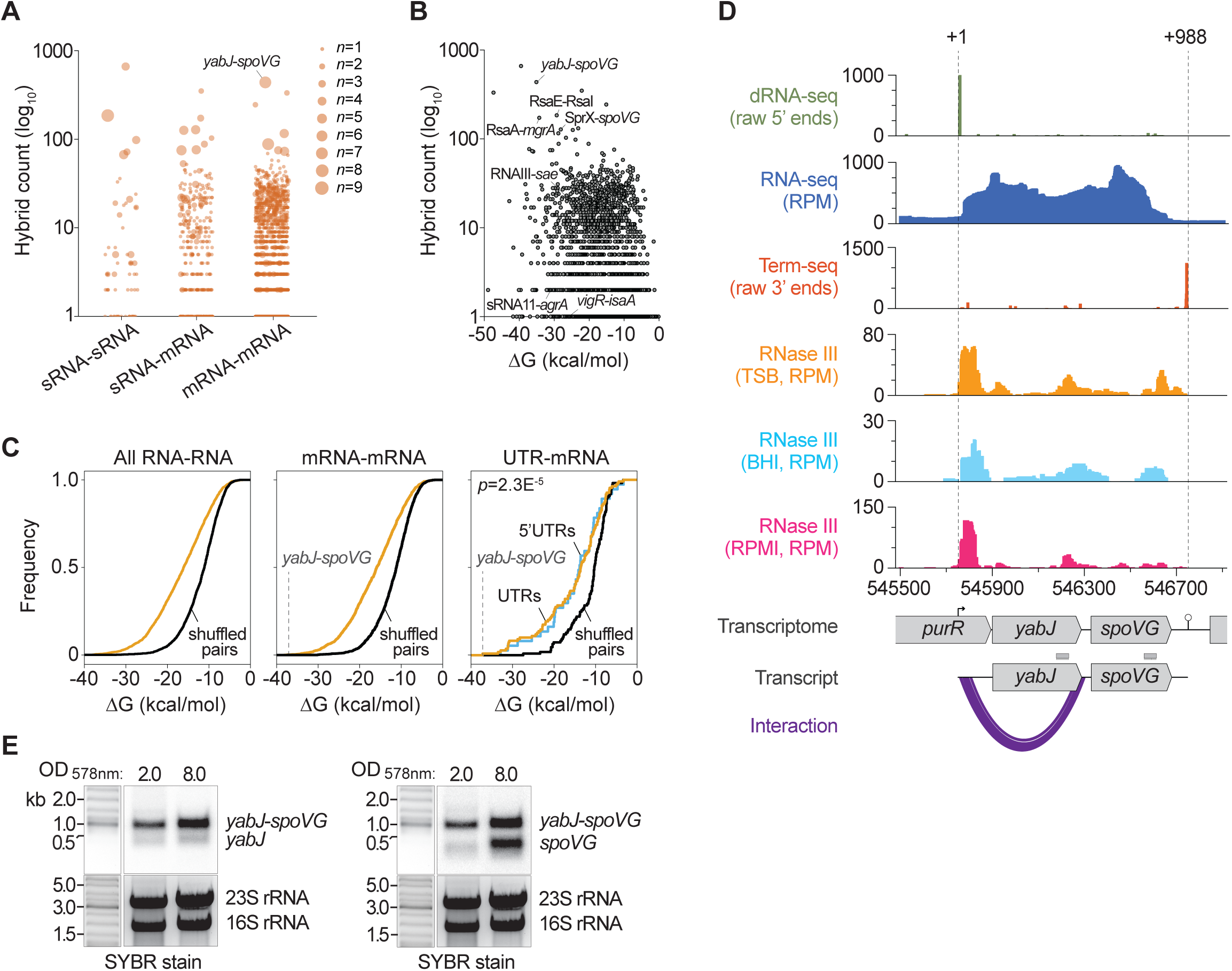
The *yabJ*-*spoVG* biscistronic transcript forms an mRNA-mRNA interaction captured by RNase III-CLASH. (**A**) Total number of RNA hybrids (*left*) distributed across each RNA-RNA interaction class (*bottom*) captured in the collated RNase III-CLASH interactome. The number of independent experiments containing the interaction is depicted by the data point size (*right*). The data point containing the *yabJ*-*spoVG* interaction is indicated (*n*=8). (**B**) Distribution of the number of RNA hybrids (*left*) and their respective RNA-RNA interaction strength (ΔG, kcal/mol). The data points containing the *yabJ*-*spoVG* interaction and *bona fide* RNA-RNA interactions are indicated. (**C**) Cumulative distribution function of RNA-RNA interaction strength (ΔG, kcal/mol) for all RNA-RNA (*left*), mRNA-mRNA (*middle*), or UTR-mRNA (right) interaction classes recovered. The distribution of ΔG of randomly shuffled RNA pairs is shown in black. The *yabJ*-*spoVG* interaction is indicated. (**D**) Histogram plots indicate RNA read data mapping to the *yabJ*-*spoVG* transcript in MRSA JKD6009. From *top* to *bottom*, dRNA-seq (raw 5’ RNA read count), total RNA-seq (reads per million, RPM), Term-seq (raw 3’ RNA read count), and RNase III-binding sites from UV-crosslinking and sequencing datasets in TSB, BHI, or RPMI media (reads per million, RPM). The position of the *yabJ*-*spoVG* transcript is indicated *below* with the x-axis representing the genomic coordinates in JKD6009. The transcription start site and termination site are indicated. The mRNA-mRNA interaction spanning the *yabJ* 5’UTR-intercistronic region is depicted in purple. (**E**) Northern blot analyses of the bicistronic *yabJ*-*spoVG* transcript, and monocistronic *spoVG* and *yabJ* mRNAs. Total RNA was extracted from VISA JKD6008 (wild-type) grown to mid-log phase (OD_578nm_ of 2.0) or stationary phase (OD_578nm_ of 8.0) and probed for *yabJ* (*left*) or *spoVG* (*right*) mRNA. SYBR Green stained rRNA is indicated *below* as a loading control.

To determine whether captured hybrids reflect genuine base-paired RNA-RNA interactions, we compared the ΔG distribution of all interactions and of the mRNA-mRNA subclasses, against a shuffled control set of randomly paired reads. Consistent with previous findings, the collated RNA-RNA interactions had significantly lower free energy than randomly shuffled pairs (**Figure 1C**, *left*). Additionally, mRNA-mRNA duplexes captured by RNase III-CLASH also had significantly lower ΔG than shuffled RNA pairs (**Figure 1C**, *middle*), confirming that hybrid reads represent thermodynamically stable RNA duplexes.

We next mapped the *yabJ*-*spoVG* interaction captured by RNase III-CLASH onto the transcript using dRNA-seq (5’ boundary ends), RNA-seq, and Term-seq (3’ boundary ends) data generated in MRSA JKD6009 [8]. dRNA-seq identified a primary transcription start site (TSS, +1) upstream of the *yabJ* coding-DNA sequence (CDS), and RNA-seq coverage across *yabJ* and *spoVG* verified their co-transcription as a bicistronic transcript terminating at a single transcription termination site (+988) downstream of the *spoVG* CDS (**Figure 1D**). Analyses of the 3’ termination sequence identified a structured RNA hairpin and poly-U tract ending at the +988 position, consistent with Rho-independent (instrinsic) termination (**Supplementary Figure 1B**). We did not identify any additional TSS or termination sites within the locus.

To confirm transcript architecture, we used Northern blotting to probe for the monocistronic *yabJ* and *spoVG* mRNAs (probe positions in **Figure 1D**). We confirm the approx. 1-kb transcript size for the full-length (bicistronic) *yabJ*-*spoVG* using either the *yabJ* or *spoVG* probe (**Figure 1E**, *left* and *right*, respectively), supporting our 5’ and 3’ RNA boundary end analyses. We further confirm the ∼550-nt mRNA for *yabJ* and ∼450-nt mRNA for *spoVG*, and find increased expression of the bicistronic transcript and *spoVG* mRNA, but not *yabJ* at the stationary phase of growth (OD_578nm_ = 8.0, **Figure 1E**).

These data confirm *yabJ*-*spoVG* as a bicistronic transcript that harbours an abundant, thermodynamically stable mRNA-mRNA interaction recovered by RNase III-CLASH in MRSA.

### *yabJ*-*spoVG* is processed by RNase III

RNA hybrid reads recovered by RNase III-CLASH were comprised between the unusually long *yabJ* 5’UTR (145-nt) and the intercistronic region of *yabJ*-*spoVG* encompassing part of the *yabJ* CDS (**Figure 1D** and **Supplementary Figure 1C**). Analyses of the hybrid reads using IntaRNA [35] verified a 46-nt RNA duplex with regions of up to 17-nt of perfect Watson-Crick base-pairing between hybrids (**Figure 2A**). We find that UTR-mRNA duplexes, including 5’UTR-mRNA duplexes captured by RNase III-CLASH had significantly lower ΔG than randomly shuffled RNA pairs (*p*=2.3E^−5^, **Figure 1C**, *right*), consistent with the formation of highly stable UTR-mRNA duplexes.

**Figure 2.**
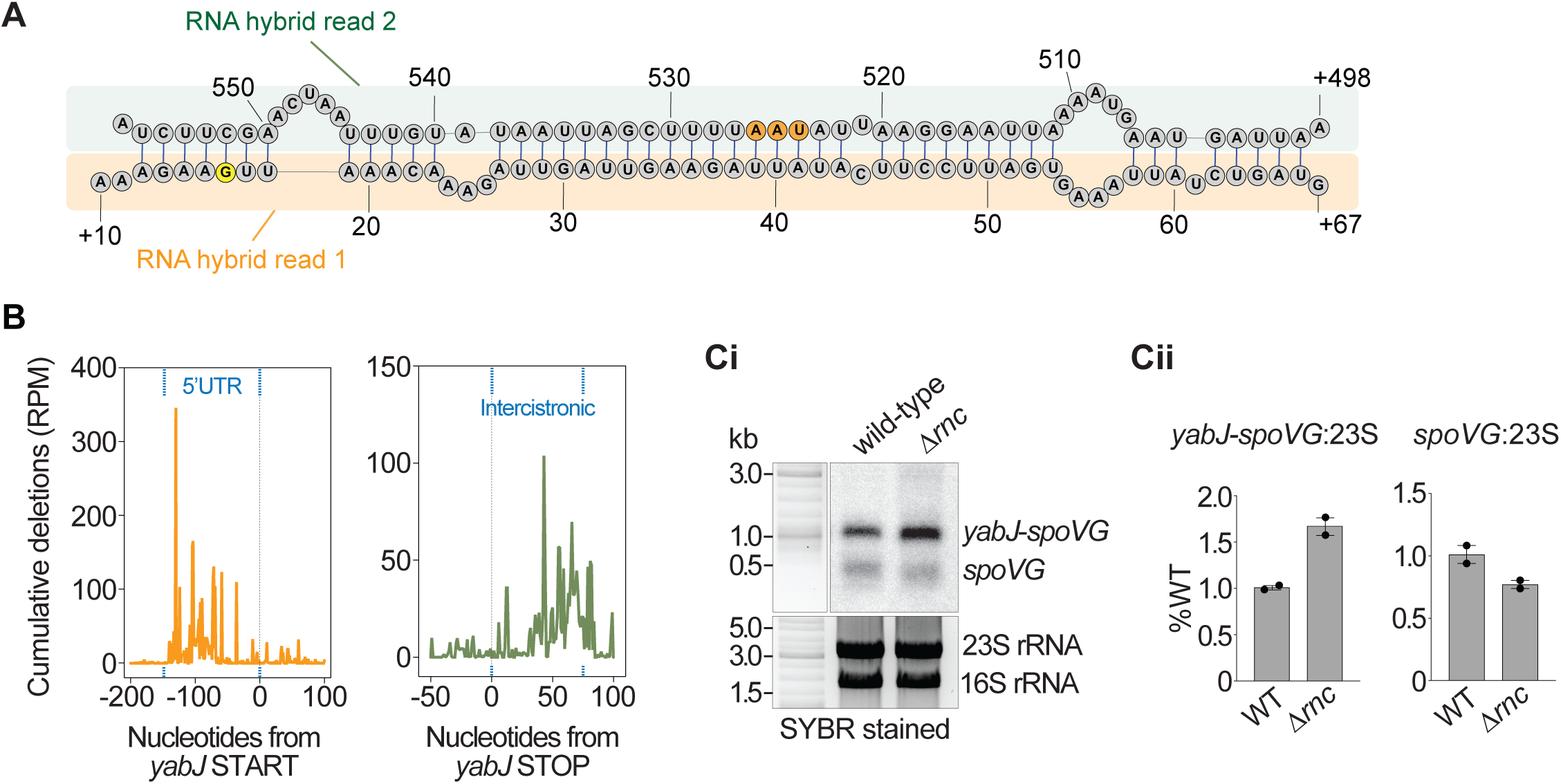
The *yabJ*-*spoVG* biscistronic transcript forms a substrate for RNase III processing. (**A**) The RNase III-CLASH identified mRNA-mRNA interaction within the *yabJ*-*spoVG* transcript at nucleotide resolution. The sequence recovered corresponding to the two hybrid reads of the RNA duplex are shown highlighted in orange (hybrid read 1) and green (hybrid read 2). The *yabJ* stop codon is indicated in orange and the guanosine with maximal RNase III contact is indicated in yellow. The nucleotide genomic positions of each hybrid read is detailed representative of the *yabJ-spoVG* transcription start site (+1 site). (**B**) RNase III contact dependent deletions mapped relative to the *yabJ* start codon (*left*) and the *yabJ* stop codon (*right*). Histogram plot represents cumulative count from UV-crosslinking and sequencing datasets in TSB, BHI, and RPMI media (reads per million, RPM). Dashed blue lines indicate the approximate position of the *yabJ* 5’UTR (*left*) and the intercistronic region (*right*). (**C**) Northern blot analyses of the bicistronic *yabJ*-*spoVG* transcript and monocistronic *spoVG* mRNA. (**i**) Total RNA was extracted from MRSA strain JKD6009 (isogenic wild-type parent) and Δ*rnc* (RNase III deletion) grown to an OD_600nm_ of 3.0 and probed for *spoVG* mRNA. SYBR Green stained rRNA are indicated below as a loading control. (**ii**) Quantification of the ratio of 23S rRNA to either the bicistronic *yabJ*-*spoVG* or monocistronic *spoVG* by densitometry, relative to wild-type.

To investigate the role of RNase III in processing *yabJ*-*spoVG*, normalised read density of RNase III-RNA binding sites in three independent growth media (TSB, BHI, and RPMI-1640 [8, 18]) were mapped to the *yabJ*-*spoVG* transcript. We find RNase III crosslinking enriched at the *yabJ* 5’UTR, consistent with the formation of a double-stranded RNA substrate for RNase III processing (**Figure 1D**). To further localise the RNase III-RNA contacts at nucleotide resolution, we mapped the point deletions introduced at UV-crosslinking sites, which mark the RNA position of protein contact. Deletions from RNase III contact were enriched within both the *yabJ* 5’UTR and the *yabJ*-*spoVG* intercistronic region (**Figure 2B**). Within the *yabJ* 5’UTR, deletions on RNase III contact was maximal at the +16 guanosine nucleotide which forms a direct base-pair with the +552 cytosine in the intercistronic region (**Figures 2A** and **2B**), consistent with RNase III cleavage favoured at GC/CG sites.

Northern analyses was used to confirm the role of RNase III in the regulation of the *yabJ*-*spoVG* transcript. We hypothesised that RNase III processing of the bicistronic transcript at the 5’UTR-intercistronic duplex would control release of monocistronic *spoVG*. Using a deletion of RNase III (*rnc*) in MRSA JKD6009, we find a 61% increase (± 4.9%) in the abundance of bicistronic *yabJ*-*spoVG* in the absence of RNase III (Δ*rnc*) relative to the isogenic wild-type (**Figure 2C**), verifying that RNase III contributes to processing and decay of the transcript. We also find a concurrent 24% decrease (± 4.6%) in abundance of the monocistronic *spoVG* in the Δ*rnc* strain relative to wild-type (**Figure 2C**), consistent with RNase III processing of *spoVG* within the bicistronic transcript.

These data demonstrate that RNase III associates with the *yabJ* 5’UTR-intercistronic duplex and processes the bicistronic transcript.

### The *yabJ* 5’UTR regulates the *spoVG* mRNA

To confirm that the *yabJ* 5’UTR forms a direct RNA-RNA interaction with the intercistronic region of *yabJ*-*spoVG,* both regions of the transcript were *in vitro* transcribed and analysed using electrophoretic mobility shift assays (**Supplementary Figure 1D**). Titration of the *yabJ* 5’UTR with radiolabelled (^32^P-labelled) *yabJ*-*spoVG* incorporating the intercistronic region shifted the ^32^P-labelled fragment to a slower migration rate consistent with the formation of an RNA-RNA complex (**Figure 3A**). The *isaA* mRNA was used as a negative control and showed no migrational shift of the ^32^P-labelled fragment (**Figure 3A**). Two competitor oligonucleotides antisense to the interaction site in either *yabJ* 5’UTR or the intercistronic region were titrated into the gel shift assay up to 100x excess concentration (**Supplementary Figure 1D**). Antisense oligomer A was able to effectively compete away at least 70.2% of the ^32^P-labelled fragment at the highest titrations, and competitor oligonucleotide B (85-mer) antisense to the *yabJ* 5’UTR competing away 91.3% of the ^32^P-labelled RNA-RNA duplex (**Supplementary Figure 1E)**, confirming a direct *in vitro* RNA-RNA interaction between the *yabJ* 5’UTR and the intercistronic region of the *yabJ*-*spoVG* transcript.

**Figure 3.**
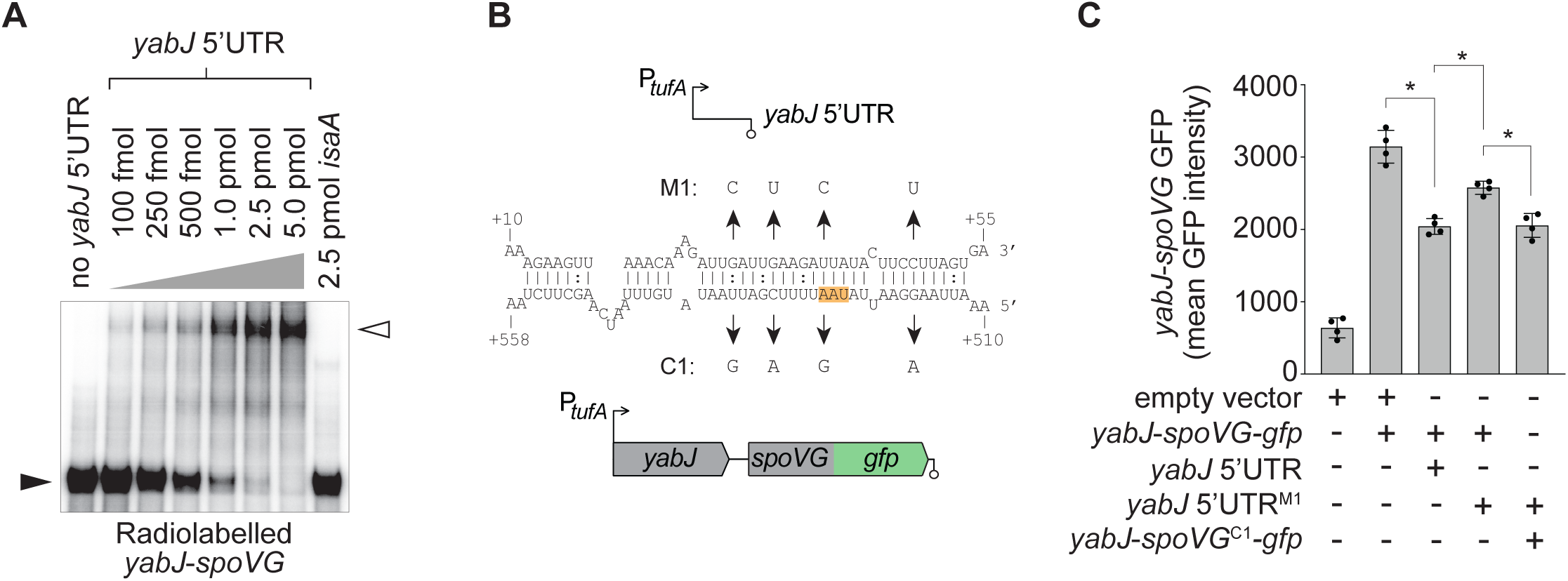
The *yabJ* 5’UTR-intercistronic region interaction represses *spoVG*. (**A**) EMSA analysis of the radiolabelled *yabJ*-*spoVG* intercistronic region (indicated as *yabJ*-*spoVG*) and the *yabJ* 5’UTR. Increasing concentrations (0 to 5 pmol, indicated *top*) of the *yabJ* 5’UTR RNA was titrated against 50 fM of radiolabelled *yabJ* 3’-*spoVG* RNA. Migration of radiolabelled RNA is indicated by the arrows. (**B**) A constitutively transcribed GFP translational fusion to SpoVG (SpoVG-GFP, indicated *bottom* panel) was expressed in *S. aureus* with or without transcription of the *yabJ* 5’UTR (indicated *top* panel). The nucleotide genomic positions of each cloned region is detailed representative of the *yabJ-spoVG* transcription start site (+1 site). The RNA-RNA interaction at nucleotide resolution is depicted with compensatory point mutations, indicated by M1 and C1, introduced into the *yabJ* 5’UTR and *yabJ*-*spoVG* intercistronic region, respectively. The *yabJ* mRNA stop codon is highlighted in orange. (**C**) The mean fluorescence intensities of constitutvely expressed construct combinations (indicated *below*) measured using a microtiter plate reader. Error bars represent standard error (*n*=4). A student’s *t*-test, two sample assuming unequal variance was used to determine statistical significance. \**p*<0.05.

Given the functional role of SpoVG as a transcription factor in *S. aureus*, we next assessed if the *cis*-regulatory RNA-RNA interaction could affect *in vivo* expression of SpoVG. The *yabJ* 5’UTR and *yabJ*-*spoVG* fused to *gfp* (encoding a YabJ-SpoVG-GFP translational fusion) were cloned into a two-plasmid system that enabled independent and constitutive expression of both transcripts, and fluorescent detection of SpoVG-GFP (**Figure 3B**). We demonstrate that expression of the *yabJ* 5’UTR was able to significantly repress SpoVG (*p*=0.00067, **Figure 3C**). Point mutations disrupting the *yabJ* 5’UTR interaction site within the region of up to 17-nt perfect Watson-Crick base-pairing (designated as *yabJ* 5’UTR^M1^, **Figure 3B**) significantly decreased repression, and repression could be restored when the complementary synonymous point mutations in *yabJ*-*spoVG* (designated as *yabJ*-*spoVG*^C1^, **Figure 3B**) were expressed together (**Figure 3C**).

Given that constitutive expression of the *yabJ* 5’UTR was able to repress SpoVG-GFP and point mutations decreased this regulatory control (**Figure 3C**), we next assessed if point mutations disrupting the *yabJ* 5’UTR interaction site on the chromosomal copy of *yabJ*-*spoVG* (designated *yabJ* 5’UTR^SNP^) would demonstrate increased expression of *spoVG* in clinical VISA isolate JKD6008 (ST239 derived from MRSA JKD6009) [36]. The *yabJ* 5’UTR^SNP^ strain was constructed by introducing point mutations at the interaction site (within the +18-48 locus position; *yabJ* start codon located at +146) (**Supplementary Figure 2A**). Consistent with the *yabJ*-*spoVG*-*gfp* fusion, Northern analyses revealed a 1.54-fold increase in bicistronic *yabJ*-*spoVG* (*p*=0.018) and a 3.57-fold increase in monocistronic *spoVG* (*p*=0.039) in the *yabJ* 5’UTR^SNP^ when compared to wild-type (**Figure 4A**). We next constructed a chromosomal repair strain using allelic exchange to restore the *yabJ* 5’UTR sequence (designated as *yabJ* 5’UTR^SNP^-repair). Using qRT-PCR with primers that anneal to either the intercistronic region of the bicistronic *yabJ*-*spoVG* (across the RNase III maximal contact site, **Figure 2B**), *yabJ*, or *spoVG*, we profiled the relative abundances of each transcript. Our qRT-PCR analyses confirmed a significant upregulation of the bicistronic transcript (*p*=1.29E^−5^) and *spoVG* mRNA (*p*=7.65E^−6^), but not *yabJ*, in the *yabJ* 5’UTR^SNP^ when compared to wild-type (**Figure 4B-D**). We also confirm restoration of both *yabJ*-*spoVG* and *spoVG* expression levels in the *yabJ* 5’UTR^SNP^-repair, demonstrating that the *yabJ* 5’UTR negatively controls the downstream *spoVG* through a *cis*-regulatory RNA-RNA interaction.

**Figure 4.**
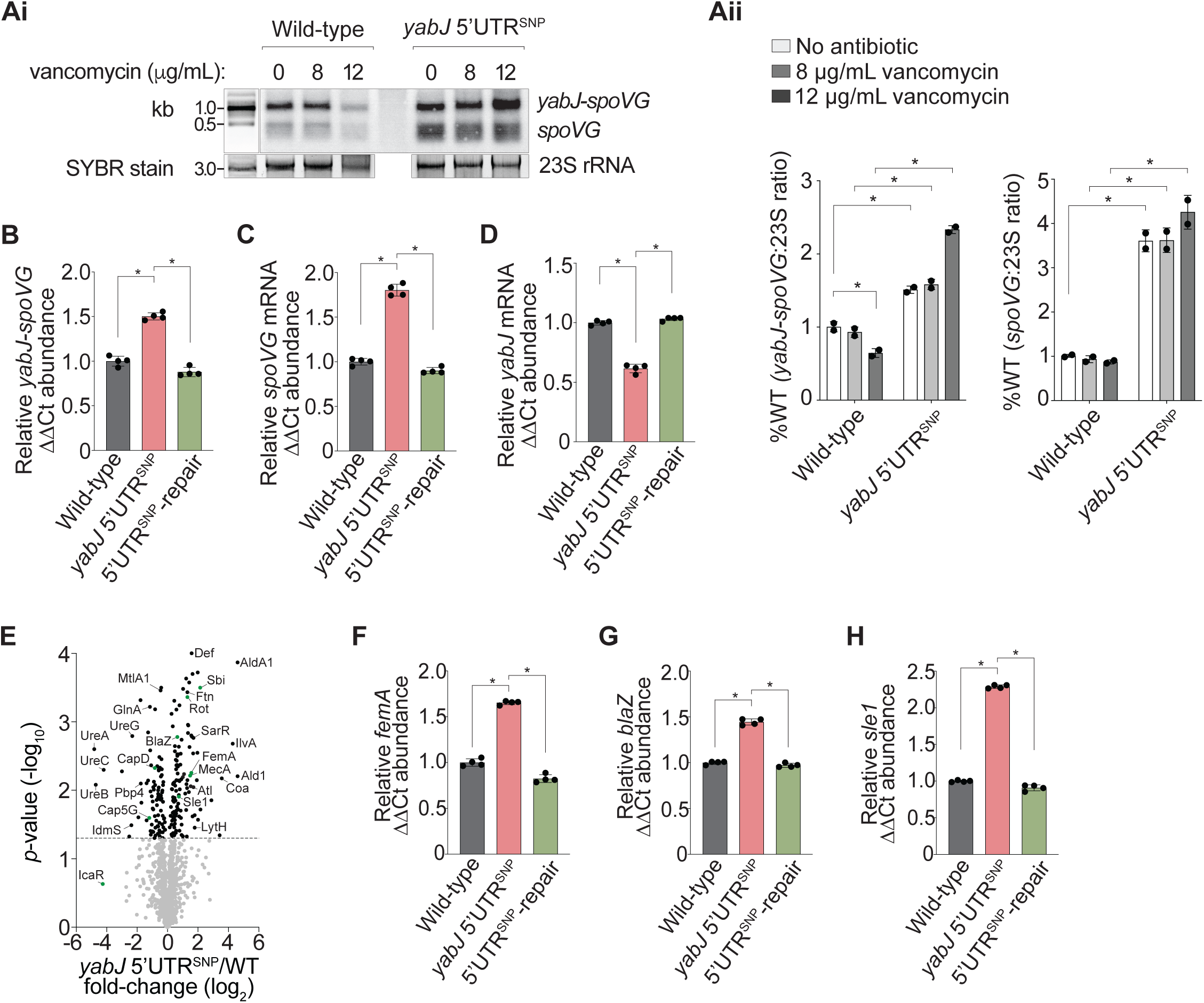
Disruption of the *yabJ* 5’UTR alleviates *spoVG* repression and coincides with changes to the SpoVG regulon. (**A**) Northern blot analyses of the bicistronic *yabJ*-*spoVG* transcript and monocistronic *spoVG* mRNA. (**i**) Total RNA was extracted from VISA strain JKD6008 (isogenic wild-type parent) and JKD6008 *yabJ* 5’UTR^SNP^ grown to an OD_600nm_ of 3.0 treated with or without increasing concentrations of vancomycin (0 to 12 μg/mL) for 1 h, and probed for *spoVG* mRNA. SYBR Green stained 23S rRNA is indicated below as a loading control. (**ii**) Quantification of the ratio of 23S rRNA to either bicistronic *yabJ*-*spoVG* or monocistronic *spoVG* by densitometry, relative to wild-type. The absence or presence of vancomycin treatment is indicated *above*. Error bars represent standard error (*n*=2). A student’s *t*-test, two sample assuming unequal variance was used to determine statistical significance. \**p*<0.05. (**B-D**) Histogram of qRT-PCR to quantify the (**B**) bicistronic *yabJ*-*spoVG* transcript, (**C**) monocistronic *spoVG*, and (**D**) monocistronic *yabJ* chromosomal expression (relative to the *gapA* mRNA) in wild-type, *yabJ* 5’UTR^SNP^, and *yabJ* 5’UTR^SNP^-repair strains. Error bars represent standard error (*n*=4). A student’s *t*-test, two sample assuming unequal variance was used to determine statistical significance. \**p*<0.05. (**E**) Volcano plot of differentially abundant proteins in the JKD6008 *yabJ* 5’UTR^SNP^ vs JKD6008 (isogenic) wild-type detected by LC-MS/MS analyses. Statistically significant changes in protein abundances are indicated in black (*p*<0.05). Known SpoVG targets are indicated in green. (**F-H**) Histogram of qRT-PCR to quantify the (**F**) *femA*, (**G**) *blaZ*, and (**H**) *sle1* chromosomal expressions (relative to the *gapA* mRNA) in wild-type, *yabJ* 5’UTR^SNP^, and *yabJ* 5’UTR^SNP^-repair strains. Error bars represent standard error (*n*=4). A student’s *t*-test, two sample assuming unequal variance was used to determine statistical significance. \**p*<0.05.

These data indicate that the *yabJ* 5’UTR directly interacts *in vivo* with the intercistronic region between *yabJ* and *spoVG* to repress *spoVG* mRNA expression.

### *yabJ* 5’UTR regulation of *spoVG* coincides with changes to the SpoVG regulon

We next assessed if the regulatory interaction between the *yabJ* 5’UTR and the *yabJ*-*spoVG* intercistronic region impacted the SpoVG regulon. The SpoVG regulon comprises targets within antibiotic resistance [24], cell wall biogenesis [26], carbohydrate metabolism [27], virulence factor production [28, 31, 32], and capsule synthesis [29]. Both wild-type and the *yabJ* 5’UTR^SNP^ strains were grown to exponential phase and total clarified protein lysates analysed using mass spectrometry (LC-MS/MS) to detect protein abundances. We identified 218 differentially abundant proteins, including 9 within the known SpoVG regulon (*p*≤0.05, **Figure 4E** and **Supplementary Table 1A**). These included genes such as *mecA* (encoding the alternative PBP2a conferring β-lactam resistance), *sbi* (encoding a secreted and surface-associated immune evasion factor), and *capD* (encoding the dehydratase required for capsular polysaccharide biosynthesis) (**Figure 4E**, highlighted in green). Ontological clustering of the significant differentially abundant proteins showed enrichment of functional roles associated with energy and amino acid metabolism, co-enzyme metabolism, transcription and translation, and cell wall biogenesis (*p*≤0.05, **Supplementary Figure 2B**), demonstrating functional overlap with the confirmed target pathways of SpoVG.

Previous work demonstrated that SpoVG directly binds and activates transcription of *femA*, *blaZ*, and *sle1*, and reported decreased expression of these targets in the Δ*spoVG* background [26]. We confirm differential abundance of FemA, BlaZ, and Sle1 in our proteomic analyses (**Figure 4E**) and demonstrate the reverse trend whereby disruption of the *yabJ* 5’UTR (*yabJ* 5’UTR^SNP^), alleviating repression of *spoVG*, led to a significantly higher abundance of these genes. We confirm these findings using qRT-PCR and demonstrate significant 1.7-fold, 1.4-fold and 2.3-fold increases in *femA*, *blaZ*, and *sle1*, respectively in the *yabJ* 5’UTR^SNP^ strain (**Figure 4F-H**), consistent with our proteomic analyses. We also confirm restoration of expression levels of the target genes in the *yabJ* 5’UTR^SNP^-repair (**Figure 4F-H**), demonstrating that the regulatory control of the *yabJ* 5’UTR on *spoVG* modulates the SpoVG regulon.

Some of the largest differentially abundant hits within our proteomic analyses included proteins not yet confirmed as targets of SpoVG, but may form part of an extended regulon. These genes include *pbp4* (encoding a cell wall penicillin-binding protein), *coa* (encoding a blood coagulase), and *ureABC* (encoding urease for the breakdown of urea). Given that UreABC was detected amongst the most negatively abundant proteins in our analyses, we assessed if the upstream *ureA* could be a novel target for SpoVG. Using qRT-PCR, we confirm a significant 3.6-fold decrease in *ureA* in the *yabJ* 5’UTR^SNP^ (*p*=0.016) and demonstrate restoration in the 5’UTR^SNP^-repair strain (**Supplementary Figure 2C**). To determine a potential SpoVG DNA-binding motif for *ureA*, genomic sequences of the general promoter region of 11 known targets within the SpoVG regulon were extracted from MRSA JKD6009 and used as input for GLAM2 software alignment (**Supplementary Table 1B**). Consensus motifs obtained identified a T_T/A_ATT_T/A_ motif enriched in SpoVG targets and this motif was used as input for GLAM2SCAN to identify the presence of a matching motif in the *ureA* promoter region (**Supplementary Figure 2D**). Two consecutive TAATT_T/A_ motifs located at the −72 and −64 sites in *ureA* were identified (**Supplementary Figure 2D**), suggesting that *ureA* and the metabolism of urea may also be under the control of SpoVG within *S. aureus*.

Collectively, disrupting the *yabJ* 5’UTR-intercistronic interaction de-represses *spoVG* and produces corresponding changes in abundance of known SpoVG regulon targets.

### *yabJ* 5’UTR controls resistance to cell wall-targeting antimicrobials

SpoVG has a broad regulon controlling genes with roles in cell wall metabolism and resistance to cell wall-targeting antibiotics [26]. Our findings demonstrate that the bicistronic *yabJ*-*spoVG* transcript and the *spoVG* mRNA (but not *yabJ*) are upregulated in late stationary phase culture conditions. Surprisingly, during increasing concentrations of acute vancomycin treatment, we find a progressive decrease in the abundance of the bicistronic *yabJ*-*spoVG* and monocistronic mRNAs (**Figure 4A** and **Supplementary Figure 3A-B**), suggesting a negative regulatory role in the acute vancomycin stress response. Notably, in the presence of increasing concentrations of vancomycin, we report significantly higher abundances of the bicistronic transcript and monocistronic *spoVG* mRNA in the *yabJ* 5’UTR^SNP^ strain when compared to wild-type (**Figure 4A**), demonstrating control of *spoVG* by the *yabJ* 5’UTR during cell wall-targeting antibiotic stress.

To further explore the functional role of the *cis*-encoded regulatory interaction to antimicrobials, we performed spot dilution assays to assess sensitivity in the presence of the cell wall-targeting β-lactam oxacillin, and the glycopeptides vancomycin and teicoplanin. In the absence of antibiotics, the *yabJ* 5’UTR^SNP^ and *yabJ* 5’UTR^SNP^-repair strains had comparable growth to VISA (isogenic wild-type) and MRSA (**Figure 5A**). However, in the presence of sub-inhibitory concentrations of oxacillin, vancomycin, and teicoplanin, growth of the *yabJ* 5’UTR^SNP^ strain was reduced 10-fold, 100-fold, and 100-fold, respectively (**Figure 5A**). We confirm sensitivity of MRSA in the presence of sub-inhibitory concentrations of both glycopeptide antibiotics and demonstrate restoration of sensitivity in the *yabJ* 5’UTR^SNP^-repair for all antibiotics assessed (**Figure 5A**).

**Figure 5.**
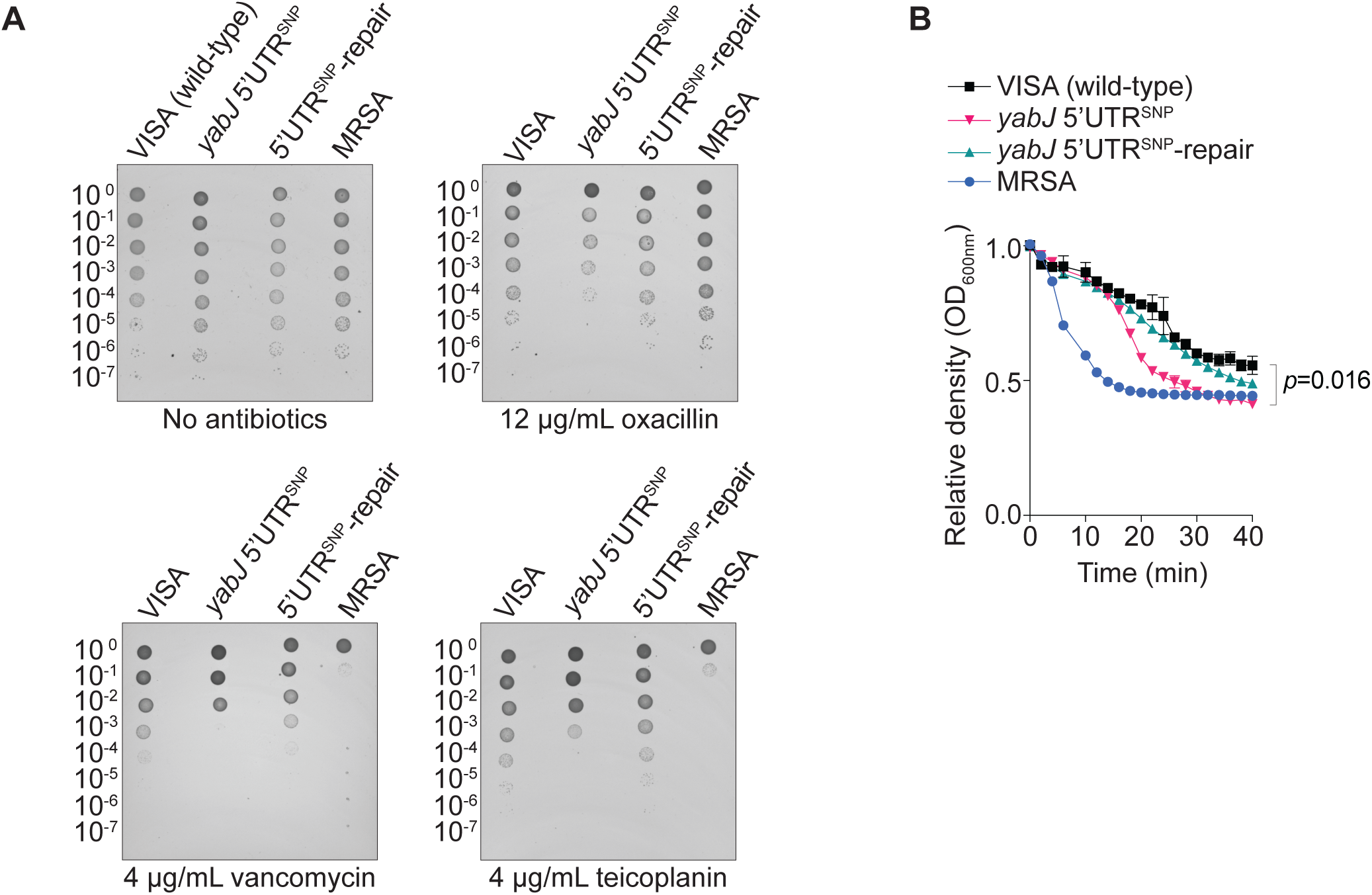
*yabJ* 5’UTR repression of *spoVG* mediates resistance to cell wall-targeting antimicrobials. (**A**) Spot dilution assays quantified in VISA JKD6008 (isogenic wild-type), *yabJ* 5’UTR^SNP^, *yabJ* 5’UTR^SNP^-repair, and MRSA JKD6009 strains in the absence (*top left*) or presence of sub-inhibitory concentrations of oxacillin (*top right*), vancomycin (*bottom left*) and teicoplanin (*bottom right*) antibiotics. Culture dilution is indicated *left*. (**B**) Lysis assay of normalised JKD6008 (wild-type), *yabJ* 5’UTR^SNP^, *yabJ* 5’UTR^SNP^-repair, and JKD6009 in MH media supplemented with 5 μg/mL lysostaphin. Error bars represent standard error (*n*=3). A student’s *t*-test, two sample assuming unequal variance was used to determine statistical significance. \**p*<0.05.

We next assessed if the *yabJ* 5’UTR^SNP^ had altered susceptibility to the cell wall-targeting antimicrobial lysostaphin that cleaves the pentaglycine crosslinks in cell wall peptidoglycan and has been explored clinically as an alternative antimicrobial agent [37]. Susceptibility to lysostaphin-mediated lysis is governed by the extent of peptidoglycan crosslinking, providing a structure-specific readout of cell wall integrity. VISA, *yabJ* 5’UTR^SNP^, *yabJ* 5’UTR^SNP^-repair, and MRSA were incubated with 5 μg/mL of lysostaphin and culture density measurements were acquired over time. The *yabJ* 5’UTR^SNP^ was significantly more sensitive to lysostaphin treatment over 40 min relative to the wild-type (*p*=0.016) and resistance was restored in the *yabJ* 5’UTR^SNP^-repair (*p*=5.01E^−5^) (**Figure 5B**). We also confirm sensitivity of MRSA to lysostaphin which had a similar bacterial density to the *yabJ* 5’UTR^SNP^ strain after 40 min of treatment (**Figure 5B**).

These data demonstrate that *yabJ* 5’UTR-mediated repression of *spoVG* is required for cell wall-targeting antimicrobial resistance in clinical VISA.

### Inducible overexpression of SpoVG phenocopies *yabJ* 5’UTR^SNP^ and attenuates cell wall thickening and biofilm formation

To understand susceptibility of the *yabJ* 5’UTR^SNP^ strain to cell wall-targeting antimicrobials and determine the impact of *spoVG* regulation on cell wall metabolism, a SpoVG overexpression construct was made using the inducible pRAB11 vector containing the *tetR*-regulated P_xyl/tet_ promoter [38] (pRAB11-SpoVG) and transformed into VISA JKD6008. Using qRT-PCR, we confirm a significant 7.4-fold increase in expression of *spoVG* at log growth phase relative to the empty pRAB11 vector (*p*<1.0E^−5^, **Supplementary Figure 3C**), verifying overexpression of our construct.

Cell wall thickening is a common phenotype of clinical VISA isolates and a key physiological differentiator to MRSA [3]. We used transmission electron microscopy (TEM) to quantify the cell wall thickness of VISA, *yabJ* 5’UTR^SNP^, *yabJ* 5’UTR^SNP^-repair, MRSA, and the pRAB11 and pRAB11-SpoVG constructs after anhydrotetracycline induction. Colonies were grown on blood agar for 16 h and fixed in gluteraldehyde solution supplemented with ruthenium red for TEM analyses. We confirm the increased cell wall thickness reported for VISA isolates relative to MRSA (*p*<1.0E^−5^, **Figure 6A**) [5, 34]. We find that the cell wall thickness in *yabJ* 5’UTR^SNP^ is significantly reduced relative to VISA (isogenic parent) (25.64 nm c.f. 39.14 nm, *p*<1.0E^−5^) and is restored to wild-type in the *yabJ* 5’UTR^SNP^-repair (40.20 nm, *p*<1.0E^−5^) (**Figure 6A** and **Supplementary Figure 3D**), demonstrating that the 5’UTR of *yabJ* contributes to cell wall thickening in VISA. Most notably, we also demonstrate a significantly reduced cell wall thickness in the pRAB11-SpoVG construct relative to the empty vector (24.96 nm c.f. 41.48 nm, *p*<1.0E^−5^) that is comparable to the *yabJ* 5’UTR^SNP^ strain (**Figure 6A** and **Supplementary Figure 3D**), indicating that regulation of *spoVG* by the *yabJ* 5’UTR is responsible for cell wall thickening in VISA.

**Figure 6.**
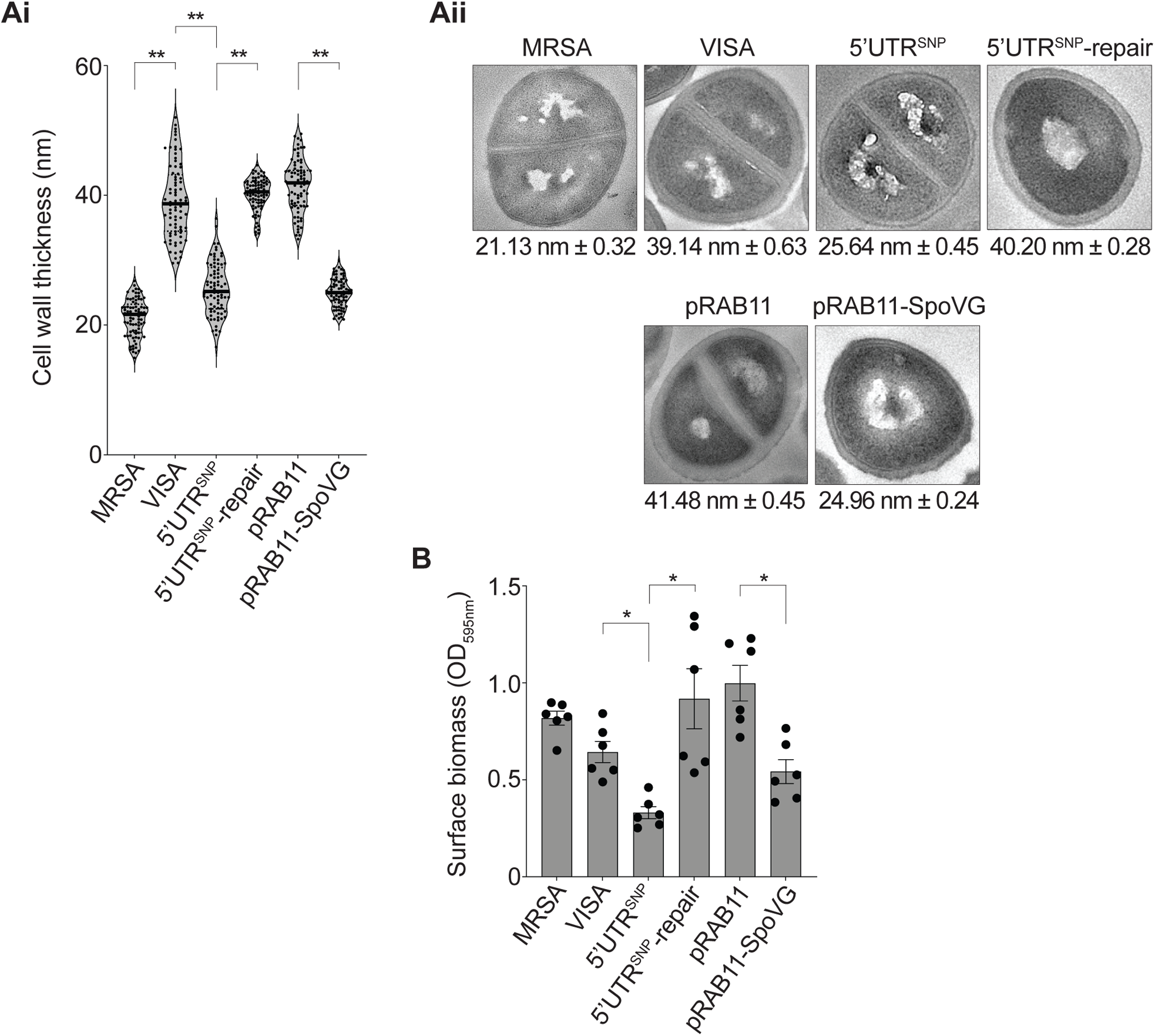
Alleviation of *spoVG* repression reverses cell wall thickening and attenuates biofilm formation in VISA. (**A**) Violin plot of cell wall thickness for MRSA JKD6009, VISA JKD6008 (isogenic wild-type), *yabJ* 5’UTR^SNP^, *yabJ* 5’UTR^SNP^-repair, pRAB11, and pRAB11-SpoVG strains (indicated *below*) as determined by transmission electron microscopy (**i**). Error bars represent standard error (*n*=80). A student’s *t*-test, two sample assuming unequal variance was used to determine statistical significance. \*\**p*<0.005. Representative transmission electron microscopy (TEM) images of MRSA JKD6009 and VISA JKD6008 derivatives (**ii**). The average cell wall thickness and standard error is shown below. (**B**) Histogram of crystal violet stained surface-associated biofilm of MRSA JKD6009 and VISA JKD6008 derivatives at 48 h of stationary growth. Error bars represent standard error (*n*=6). A student’s *t*-test, two sample assuming unequal variance was used to determine statistical significance. \**p*<0.05.

Previous work has shown that deletion of *spoVG* significantly increased biofilm development in a foodborne *S. aureus* isolate [30]. We hypothesised that alleviating *spoVG* repression would lead to reduced biofilm development in VISA. To assess if *yabJ* 5’UTR regulation of *spoVG* modulated biofilm formation, we quantified the attached crystal violet stained biomass for each normalised culture after 48 h incubation. We find an increased biofilm formation in MRSA relative to VISA (*p*=0.027, **Figure 6B**), consistent with previous observations of diminished pathogenesis in VISA isolates [3]. Notably, biofilm development was significantly reduced in the *yabJ* 5’UTR^SNP^ strain relative to VISA (*p*=0.0011) and restored in the *yabJ* 5’UTR^SNP^-repair strain (*p*=0.012, **Figure 6B**). Importantly, we also demonstrate significantly reduced biolfilm development in the pRAB11-SpoVG construct relative to the empty pRAB11 vector (*p*=0.0028), demonstrating that the *yabJ* 5’UTR biofilm development phenotype is consistent with overexpression of SpoVG.

Collectively, these results indicate that overexpression of SpoVG phenocopies the *yabJ* 5’UTR^SNP^ and demonstrates that *yabJ* 5’UTR-mediated de-repression of *spoVG* attenuates the cell wall thickening process and biofilm development in VISA.

## DISCUSSION

Bacterial genes are often arranged in polycistronic operons allowing rapid and coordinated transcription. Both transcript stability [14, 39, 40] and translation initiation factors [41, 42] control relative gene expression, however post-transcriptional gene regulation within co-transcribed polycistronic units remain incompletely understood. In this study, we identified a *cis*-regulatory RNA-RNA interaction between the 5’UTR of *yabJ* and the intercistronic region of the *yabJ*-*spoVG* bicistronic transcript. We confirm that this *cis*-encoded interaction occurs through direct base-pairing to negatively regulate the monocistronic *spoVG* mRNA. Disruption of the *cis*-interaction re-sensitised a clinical VISA isolate to cell wall-targeting antimicrobials, including the clinically-relevant glycopeptide antibiotics used to treat MRSA infections. We demonstrate that alleviation of *spoVG* repression from the *yabJ* 5’UTR coincided with changes to the SpoVG regulon, and attenuated cell wall thickening and biofilm development in VISA, likely contributing to the increased sensitivity to cell wall-targeting antimicrobials.

Several examples of *cis*-acting regulation exist in prokaryotes, however to our knowledge, this is the first mRNA-mRNA interaction within a polycistronic operon that has *cis*-encoded regulatory functions. Our 5’ and 3’ RNA boundary end data confirms that the *yabJ* 5’UTR sequence is not independently transcribed or prematurely terminated, indicating that the regulatory function is intrinsic to the mRNA 5’UTR. The long 3’UTR of *hilD* in *Salmonella enterica* negatively regulates the *hilD* transcript likely through the dual combination of RNase E- and PNPase-mediated degradation [43]. In *S. aureus*, the *icaR* mRNA (encoding a repressor of biofilm development) forms a *cis*-regulatory interaction at the 5’ and 3’UTRs generating a substrate for RNase III processing. Disruption of this *cis*-encoded interaction increased expression of *icaR* and attenuated biofilm formation [21]. We report extention of this RNase-dependent *cis*-regulatory theme, demonstrating that the bicistronic *yabJ*-*spoVG* transcript forms a substrate for RNase III-mediated endonucleolytic processing and partial generation of monocistronic *spoVG*.

The *yabJ*-*spoVG* interaction is the most abundant mRNA-mRNA duplex associated with RNase III in our interactome where we confirmed RNase III-mediated regulation of the *vigR*-*dapE* and *vigR*-*isaA* duplexes in MRSA [8, 20]. Deletion of RNase III increased bicistronic *yabJ*-*spoVG* and decreased monocistronic *spoVG*, and RNase III-RNA binding sites and contact deletions were enriched within the *yabJ* 5’UTR, consistent with RNase III processing an RNA duplex at the 5’UTR-intercistronic region of the transcript. The site of maximal RNase III contact (the +16 GC base-pair) sits within this duplex upstream of the *spoVG* translation initiation region. RNase III cleavage at this site would likely release an intact monocistronic *spoVG* that remains available for translation. The modest reduction in monocistronic *spoVG* release in the absence of RNase III indicates possible endonucleolytic redundancies. This is most evident in *E. coli* where the antisense-encoded sRNA GadY directs cleavage of the intercistronic region of the bicistronic *gadXW* transcript that is only partially RNase III-mediated with a likely dual requirement for RNase E to allow release of stable monocistronic *gadX* and *gadW* mRNA [16, 44, 45]. RNase E is absent in *S. aureus*, although the exoribonuclease RNase J1 contains partial endonucleolytic activity, and notably, suppressor mutations in RNase J1 restored a non-motile phenotype in the Δ*spoVG* background of *Listeria monocytogenes* [33], suggesting a potential link between *spoVG* and RNase J1 regulation.

The 5’UTR of *yabJ* is unusually long at 145-nt, while the average 5’UTR length for mRNA transcripts in *S. aureus* is 41-nt [8]. Long UTRs within Gram-positive *Bacillota* have emerged as large post-transcriptional regulatory hubs controlling transcript stability and translation [19, 46–48]. Additionally, intercistronic regions are often targeted by broad mechanisms of regulation [14, 16, 49]. Disruption of the RNA-RNA interaction seed within *yabJ-spoVG* alleviated repression of *spoVG* on the chromosome and from a translational fusion construct. Using proteomics and qRT-PCR, we demonstrated modest but significant changes to the SpoVG regulon, including targets within antibiotic resistance (notably *blaZ*, encoding β-lactamase) and cell wall peptidoglycan turnover (notably *femA* and *sle1*, encoding an aminoacyltransferase and autolysin, respectively) [26]. Beyond these established targets, we independently validated *ureA* (encoding the γ-subunit of urease) using qRT-PCR and motif analyses as a likely additional target within the SpoVG regulon. Urease degradation and nitrogen metabolism have been implicated in the pathogenesis of *S. aureus* [50, 51], and we find enrichment of energy and amino acid metabolism in our ontological clustering analyses. The UreABC and remaining differentially abundant proteins detected in our proteomic analyses will require further confirmation in future work.

Intermediate vancomycin resistance is the most common cause of treatment failure in pateints with MRSA infections [3]. Isolates of MRSA that have acquired intermediate resistance are commonly associated with a thicker cell wall. De-repression of *spoVG* reversed cell wall thickening in VISA and an inducible SpoVG overexpression construct phenocopied this reduction in cell wall thickness, demonstrating causation of SpoVG in the cell wall thickening pathway. Interestingly, we find that the bicistronic and monocistronic transcripts are repressed during acute vancomycin treatment in VISA, suggesting that repression of SpoVG forms part of an adaptive stress response to limit the pool of nascent D-alanyl-D-alanine peptidoglycan residues at mid-cell for glycopeptide binding [52]. This is congruent with observations in *L. monocytogenes*, whereby deletion of *spoVG* increased resistance to the cell wall-targeting antimicrobial lysozyme which hydrolyses the glycosidic bonds between the peptidoglycan sugar monomers [33]. Disruption of the *yabJ* 5’UTR alleviates *spoVG* repression and re-sensitises VISA to cell wall-targeting antimicrobials likely through the combined loss of a thick cell wall allowing antimicrobial uptake and diffusion, and the inability to modulate nascent peptidoglycan metabolism and availability.

In *S. aureus*, deletion of *spoVG* increased biofilm formation *in vitro* [30] and bacterial burden in a murine skin infection model [32], and in *L. monocytogenes* produced a hypervirulent phenotype [33]. Consistent with these findings, we demonstrate *yabJ* 5’UTR-mediated de-repression of *spoVG* produced the reverse trend, leading to significantly reduced biofilm development in VISA. This is possibly independent of cell wall thickness and likely mediated through down-regulation of the virulence-associated pathways in the regulon, including repression of cell aggregation factors [31], quorum-sensing systems [32], and exotoxin secretions [32].

The growing recognition of regulatory UTR-mediated mRNA-mRNA interactions indicate that regulatory mRNA are a widespread strategy for coordinating gene expression in Gram-positive *Bacillota* [19]. Post-transcriptional, sub-operonic gene regulation provides an advanced framework for selectively controlling individual genes within polycistronic systems. This aligns with renewed interest in RNA-based synthetic biology approaches, where engineered regulatory RNA elements are increasingly used to modulate transcript stability, processing, and translation, enabling predictable tuning of gene-specific expression [53]. *Cis*-acting elements of this kind expand the genetic toolbox for rewiring bacterial gene expression from within, rather than in *trans* of the operons that encode it.

## METHODS

### Bacterial strains, plasmids and general culture conditions

*S. aureus* was routinely cultured at 37°C on solid or in liquid brain heart infusion (BHI) media unless otherwise specified. The complete genome sequences for MRSA JKD6009 [34] and VISA JKD6008 [36] have been reported previously. *E. coli* strains were cultured at 37°C on solid or in liquid Luria-Bertani (LB) media. Antibiotics were routinely used to select for plasmids in *S. aureus* at 15 μg/mL chloramphenicol and in *E. coli* at 100 μg/mL ampicillin, unless otherwise specified. All bacterial strains were stored at −80°C as stationary phase cultures with 16% (v/v) glycerol (**Supplementary Table 2A)**.

### Strain modifications

The *S. aureus yabJ* 5’UTR^SNP^ and *yabJ* 5’UTR^SNP^-repair strains were constructed using the previously described pIMAY-Z vector and allelic exchange system [54]. Briefly, at least 500-nt flanking regions from VISA JKD6008 were synthesised as double-stranded DNA fragments (IDT) (**Supplementary Table 2B**) and amplified using Phusion DNA polymerase (NEB). Transformants were passaged and selected on solid BHI and confirmed using allele-specific PCR and genomic sequencing. Loss of pIMAY-Z was confirmed by chloramphenicol sensitivity and plasmid-specific PCR (**Supplementary Table 2B**). The construction of pRAB11-SpoVG was performed by amplifying *spoVG* (CDS) from VISA JKD6008 using Phusion DNA polymerase (NEB) with primers containing the KpnI and KasI restriction sites, and the *rrn1* T7 terminator (**Supplementary Table 2C**). The amplified product was cloned into cut pRAB11 [38] and colonies screened using PCR with primers flanking the insertion site (**Supplementary Table 2B**). The pRAB11-SpoVG construct was confirmed by whole plasmid sequencing, transformed into electrocompetent *E. coli* IM08B and then transformed into electrocompetent JKD6008.

### Interactome analysis

The pipeline for identifying RNA-RNA and RNA-protein interactions from CLASH datasets has been described previously [8, 18] and implemented in a Snakemake workflow available in GitHub (https://github.com/IgnatiusPang/Hyb-CRAC-R) [55]. The identification of the *yabJ-spoVG* transcript that formed the sequence hybrid reads representing the RNA-RNA interaction was performed using Hyb software [56]. The counts corresponding to each RNA from the hybrid reads was identified using pyCRAC software [57]. Statistical analysis of the RNA-RNA interactions was calculated using previously described R scripts [55, 58] and the free energy of hybridisation determined using RNAduplex [59]. Sequence features including 5’ and 3’UTR annotations were obtained from dRNA-seq and Term-seq datasets [8]. RNA hybrid reads were used as input into the IntaRNA (ver5.0) [35] and RNAstructure (ver6.6) [60] software programs for *in silico* confirmation of the *yabJ*-*spoVG* interaction. High confidence sites of direct RNA contact with RNase III were identified using contact-dependent deletions as described previously [61]. Independent prediction of Rho-independent (intrinsic) termination was performed using TerminatorNET software [62].

### Translational GFP fusions

The *yabJ*-*spoVG* transcript was amplified from VISA JKD6008 using Phusion DNA polymerase (NEB) and cloned into the pCN33 vector [63] at the BglII/EcoRV enzyme sites to allow translational read-through to *gfp* under constitutive expression of the P*_tufA_* promoter (**Supplementary Table 2C**). The *yabJ* 5’UTR sequence was amplified from VISA JKD6008 using Phusion DNA polymerase (NEB) and cloned into the pICS3 vector [63] at the PstI/EcoRI enzyme sites, allowing co-constitutive expression with pCN33 as previously described [8]. Plasmid constructs were verified by PCR using primers flanking the insert region and Sanger sequencing. Point mutations were introduced using the QuikChange II site-directed mutagenesis kit (Agilent Technologies) according to the manufacturer’s instructions (**Supplementary Table 2C**) and confirmed using Sanger sequencing. Plasmid constructs were co-transformed into *S. aureus* strain RN4220 [64] and selected at 24 h of growth at 37°C on solid BHI supplemented with 10 μg/mL erythromycin and 10 μg/mL chloramphenicol. Co-transformed RN4220 strains were stored at −80°C as stationary phase cultures with 16% (v/v) glycerol. Individual colonies grown on BHI supplemented with 10 μg/mL erythromycin and 10 μg/mL chloramphenicol were used to inoculate 1 mL of 0.45 μm filtered liquid BHI and grown 16 h at 37°C with 200 rpm shaking. Cultures were then diluted to an OD_600nm_ 1.0 into 0.45 μm filtered PBS (pH 7.4) and aliquoted into black 96-well microtitre plates (Thermo). The mean cellular fluorescence intensity for each culture was quantified (*n*=4) using a Tecan Spark microplate reader at the default GFP wavelength setting and a student’s *t*-test, two-sample assuming unequal variance was used to determine significance.

### Northern blot

Total RNA was extracted and purified from *S. aureus* using the GTC-phenol:chloroform method as previously described [65]. At least 5 μg of total RNA was denatured with glyoxal containing bromothymol blue and xylene cyanol at 55°C for 1 hr. Denatured RNA was resolved on a 1% BPTE-agarose gel containing SYBR Green (Thermo) and run at 100 V in 1x BPTE buffer for 1 hr. Intact ribosomal (r)RNA was confirmed and visualised on a Bio-Rad Chemi-doc imager. The gel was washed consecutively in 200 mL of 75 mM NaOH, 200 mL of neutralisation solution (1.5 M Nal and 500 mM Tris-HCl, pH7.5), and 200 mL of SSC buffer (3 M NaCl and 300 mM sodium citrate, pH 7.0) for 20 min each. RNA was capillary transferred onto a hybond-N+ nylon membrane (GE Healthcare) and UV-crosslinked in a Stratagene auto-crosslinker with 1200 mJ UV-C. The membrane was equilibrated in Ambion ULTRAhyb hybridisation buffer at 42°C for at least 1 h and then incubated with 10 pMol of 30 μCi γ^32^P-ATP-labelled oligonucleotide probe (**Supplementary Table 2B**) at 42°C for 16 h. Membranes were washed in SSPE (sodium chloride sodium phosphate EDTA) buffer with the addition of 0.1% SDS at 42°C for 3x 15 mins and imaged using a BAS-MP 2040 phosphorscreen on a FLA9500 Typhoon (GE Healthcare).

### qRT-PCR

VISA JKD6008 and derivative cultures were diluted 1:100 into 10 mL fresh liquid BHI and grown at 37°C with 200 rpm shaking to OD_600nm_ 3.0. Cultures containing pRAB11 were supplemented with 100 ng/mL anhydrotetracycline and 15 μg/mL chloramphenicol. Cells were harvested by spinning at 5,000 *g* for 10 min at 4°C. A total of 5 U of RNasin (Promega) and 10 U of RQ1 RNase-free DNase (Promega) was added and RNA purified using the GTC-phenol:chloroform extraction procedure [65] with the addition of 10 μg/mL lysostaphin (Merck) to aid cell lysis. At least 1 μg of RNA was reverse-transcribed using SuperScript IV (Thermo), according to the manufacturer’s instructions. qPCR was performed on a Bio-Rad CFX Opus real-time thermocycler using SensiFAST SYBR Hi-ROX mastermix (Bioline), according to the manufacturer’s instructions. A total cDNA concentration of 100 ng in combination with 400 nM oligonucleotides per reaction (**Supplementary Table 2B**) resulted in ideal Ct values of between 8-12 for relevant controls. Relative gene expression was determined using ΔΔCt abundance of the *gapA* (SAA6008_RS08745, glyceraldehyde-3-phosphate dehydrogenase) transcript as a reference control [8, 66]. Statistical significance was determined using a two-tailed Student’s *t*-test assuming unequal variances. Data presented as mean ± SEM.

### Electrophoretic mobility shift assay

The *yabJ* 5’UTR and *yabJ*-*spoVG* transcript (encompassing the 3’ transcript end of *yabJ* mRNA and the *spoVG* mRNA) were *in vitro* transcribed (IVT) using 40 U of T7 RNA polymerase (Roche) (**Supplementary Table 2B**). IVT products were RQ1 DNase treated (Promega) for 15 mins at 37°C, phenol-chloroform extracted and ethanol precipitated, and then separated on a 6% polyacrylamide TBE-6M urea gel. Products were excised, crushed, and incubated in 500 μL RNA gel elution buffer (10 mM magnesium acetate, 0.5 M ammonium acetate, 1 mM EDTA) for 16 h at 4°C. RNA was extracted from the eluate using phenol-chloroform extraction and ethanol precipitation. Approximately 50 pM of *yabJ*-*spoVG* RNA was dephosphorylated using alkaline phosphatase (Thermo), then extracted using phenol-chlorofom and ethanol precipitation. The 5’ ends of *yabJ*-*spoVG* were radiolabelled with 30 μCi γ^32^P-ATP using T4 polynucleotide kinase (NEB). The radiolabelled product was separated from free nucleotides using a MicroSpin G-50 column (Cytiva), and purified on a denaturing PAGE gel as above. Increasing excess amounts of the *yabJ* 5’UTR RNA were added to 25 fM of radiolabelled *yabJ*-*spoVG* in 1x duplex buffer. Where appropriate 0.25 μM of antisense competitor oligonucleotides (**Supplementary Table 2B**) were added to compete away radiolabelled *yabJ*-*spoVG* at a concentration excess of 100x. RNA was annealed, run and visualised as above.

### Protein extracts and LC-MS/MS

VISA strain JKD6008 (isogenic parent) and *yabJ* 5’UTR^SNP^ were grown in liquid BHI to an OD_600nm_ 2.0 in biological triplicates. Cultures were harvested by centrifugation (5,000 *g* for 10 min) and 1 mL of lysis buffer (50 mM Tris-HCl (pH 7.8), 150 mM NaCl, 0.1% NP-40 and 1 tablet of “cOmplete” EDTA-free protease inhibitor (Roche) per 50 mL of buffer) and 2 V of 0.1 mm zirconia beads added to each cell pellet and vortexed for 2 x 40 sec intervals using a FastPrep-24 5G (MP Biomedicals). Dry ice was used to ensure cell pellets stayed chilled during intervals. Cell debris was centrifuged (4,500 *g* for 20 min) and the clarified lysate was transferred to 1.5 mL microcentrifuge tubes and further clarified at 16,000 *g* for 20 min. Lysates containing at least 50 μg of total protein were analysed by LC-MS/MS on a Fusion LUMOS instrument (Thermo) (Mark Wainwright Analytical Centre, Sydney). Raw label free quantification (LFQ) of the data was conducted using Fragpipe (v22) [67]. Protein identification was performed against the annotated proteome of JKD6008 (NCBI GenBank CP002120) using the LFQ-MBR workflow. The output based intensities was filtered to remove contaminants and retain proteins with at least 2 non-missing values for at least 1 group. Data was normalised using variance stabilisation analyses [68]. Differential protein abundance between groups was assessed using Welch’s t-test, with Benjamini-Hochberh correction applied [69]. Data are available at the ProteomeXchange Consortium via the PRIDE partner repository with the dataset identifier PXD084383 (reviewer token: MUyL37FhA4WN).

### Motif analysis

Genomic sequences of *blaZ*, *esxA*, *femA*, *fmtB*, *icaR*, *mecA*, *rot*, *sarR*, *sbi*, *sle1*, and *spa* genes in MRSA JKD6009 encompassing the first 10 amino acids and up to 100-nt upstream of the start codon (within the 5’UTRs) were extracted (**Supplementary Table 1B**). To identify conserved sequence motifs, genomic sequences were used as input for GLAM2 alignment analyses (MEME Suite ver5.5.9) [70] with the following options: minimum aligned sequence count of 5 and 20 alignment replicates. Consensus motifs obtained from this analyses were then used as input for GLAM2Scan [70] to search for the presence of a matching motif in *ureA.* The genome viewer SnapGene was used to confirm the presence of the sequence motif within the promoter region (5’ transcriptional start site of *ureA* was determined from dRNA-seq data in MRSA JKD6009 [8]).

### Antibiotic spot assay

MRSA JKD6009, VISA JKD6008 (isogenic parent), *yabJ* 5’UTR^SNP^, and *yabJ* 5’UTR^SNP^-repair strains were used to inoculate 5 mL of liquid Meuller-Hinton (MH) and grown for 16 h at 37°C with 200 rpm shaking. Cultures were aliquoted into a 96-well microtitre plates, serially diluted (up to 10^−7^) into liquid MH and spotted onto solid MH plates supplemented with or without oxacillin, vancomycin, and teicoplanin. Spot plates were air dried at room temperature and incubated at 37°C for 24 h. Plates were imaged on a Bio-Rad Chemi-doc using the default trans-white light setting.

### Lysis assay

MRSA JKD6009, VISA JKD6008 (isogenic parent), *yabJ* 5’UTR^SNP^, and *yabJ* 5’UTR^SNP^-repair strains were used to inoculate 5 mL of liquid MH and grown for 16 h at 37°C with 200 rpm shaking. Cultures were centrifuged (500 *g* for 5 min), washed in sterile PBS and normalised to an OD_600nm_ 1.0 in liquid MH. Cultures were aliquoted into 96-well microtitre plates with the addition of 5 μg/mL lysostaphin (Merck) and allowed to incubate with gentle shaking at 37°C for 40 mins. The mean OD_600nm_ absorbance for each culture was quantified (*n*=3) at 2 min intervals using a Spark 20M multimode microplate reader (Tecan). A student’s *t*-test, two-sample assuming unequal variance was used to determine significance.

### Transmission electron microscopy

MRSA JKD6009, VISA JKD6008 (isogenic parent) and derivative cultures were cultured on solid Columbia horse blood (Thermo) at 37°C for 24 h. Strains containing pRAB11 were cultured with 100 ng/mL anhydrotetracycline and 5 μg/mL chloramphenicol. Colonies were scraped from the agar surface, resuspended in 1 mL sterile PBS, and centrifuged at 500 *g* for 5 min. Cell pellets were resuspended in 2.5% glutaraldehyde containing 0.1% ruthenium red and fixed for 24 h. Cells were pelleted by centrifugation (500 *g* for 5 min), resuspended in 0.1 M dibasic sodium phosphate buffer (Na_2_HPO_4_), and prepared for TEM as previously described [8, 34]. Cells were analysed on the Talos L120C G2 microscope and images were acquired up to 36,000 magnification. To determine cell wall thickness, two diametrically opposite measurements from 80 individual horizontally-planar cells were recorded using ImageJ. The student’s *t*-test, two-sample assuming unequal variance was used to determine statistical significance. Data presented as mean ± SEM.

### Biofilm assay

Surface-associated biofilm formation by MRSA JKD6009, VISA JKD6008, and derivative strains was quantified using a crystal violet microtiter plate assay in 96-well plates, as previously described [71] with minor modifications. Briefly, single colonies were used to inoculate 5 mL of liquid tryptic soy broth (TSB) and incubated at 37°C with 220 rpm shaking for 16 h. Strains carrying the pRAB11 vector were supplemented with 15 µg/mL chloramphenicol. Overnight cultures were normalised to an OD_600nm_ of 0.1 in fresh TSB containing 0.5% glucose, with pRAB11 cultures supplemented with 100 ng/mL anhydrotetracycline and 15 µg/mL chloramphenicol. Aliquots (100 µL) of each cell suspension were transferred to a 96-well microtiter plate and incubated statically at 37°C for 48 h. Wells were washed 3x with sterile water using an ELx405 automated plate washer (BioTek) to remove planktonic cells. Biofilms were stained with 0.1% crystal violet for 10 min at ambient temperature, washed 3x with sterile water to remove excess stain, and air-dried. Bound crystal violet was solubilised for quantitative analyses using 33% glacial acetic acid for 15 min at ambient temperature, and OD_595nm_ absorbance was measured using a Spark 20M multimode microplate reader (Tecan). Data presented is the mean ± SEM from 6 biological replicates with each biological replicate the average of 2 technical replicates (*n*=6). Statistical significance was determined using a two-tailed Student’s *t*-test assuming unequal variances.

### Statistics and reproducibility

All Northern blots and EMSA analyses were performed a minimum of twice and the images presented are representative of results from replicate experiments.

## Acknowledgements

The authors would like to thank Joyce To (Australian Institute for Microbiology and Infection) for technical assistance. R.L. was supported by PhD program scholarships from the University of Technology Sydney and RNA Australia. D.G.M. was supported by a Chancellor’s Research Fellowship from the University of Technology Sydney and a Strategic Project Grant from the New South Wales RNA Reseach & Training Network (NSW-RRTN) (RG251841). J.J.T. and D.G.M. were supported by funding from the National Health and Medical Research Council (NHMRC) (GNT2028572).

---

**Supplementary Figure 1.**
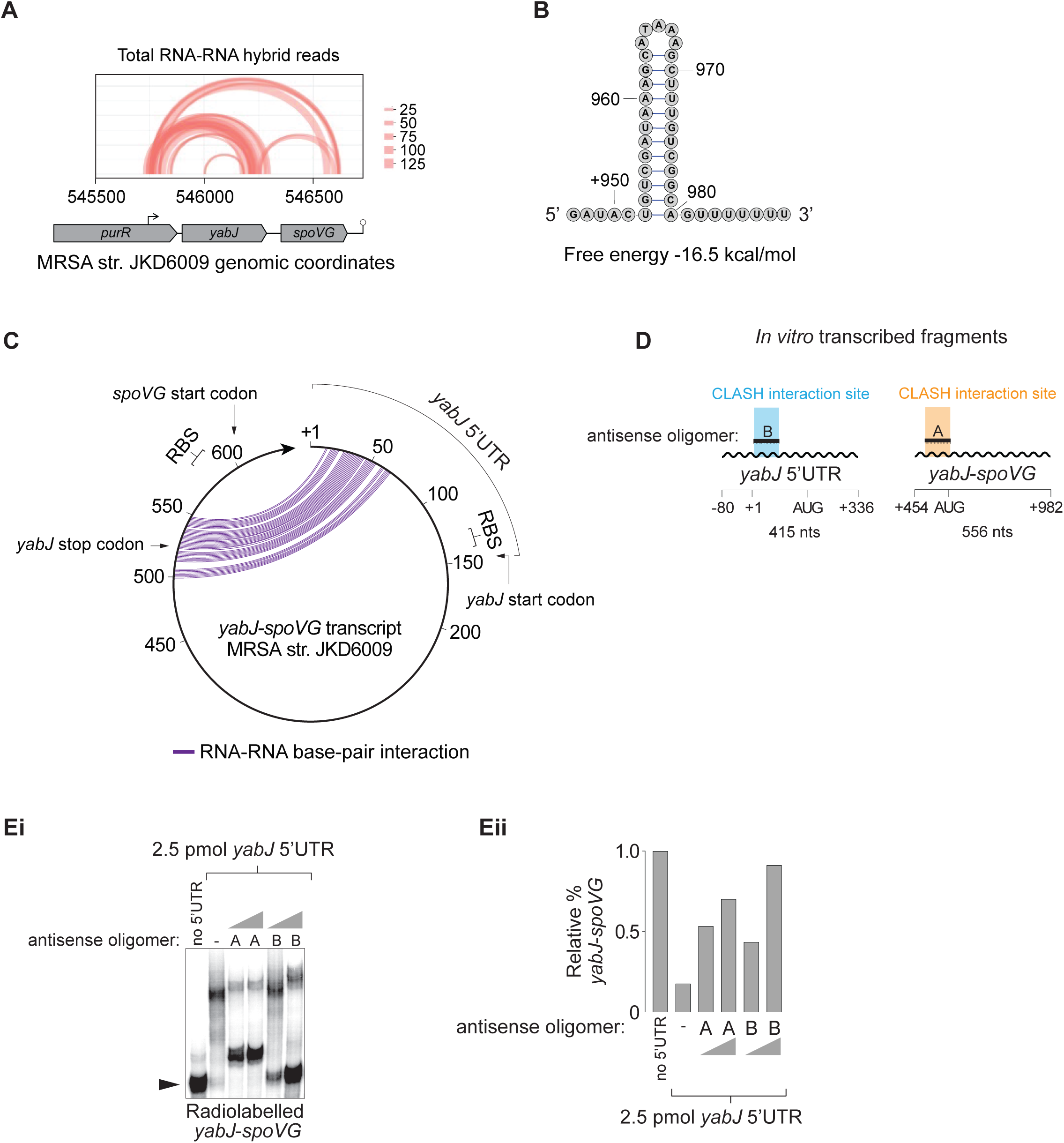
(**A**) Representation of the *yabJ*-*spoVG* bicistronic operon and mapping of total RNA-RNA hybrid reads recovered from RNase III-CLASH in MRSA strain JKD6009. Number of hybrid reads is indicated *right*. Transcript architecture of the bicistronic *yabJ*-*spoVG* is *below* (**B**) Predicted intrinsic terminator associated with *yabJ*-*spoVG* in MRSA. Genomic positions relative to the *yabJ-spoVG* transcription start site (+1 site) is indicated. (**C**) Schematic circular representation of the *yabJ*-*spoVG* transcript in the MRSA strain JKD6009. The nucleotide genomic positions of the start and stop codons, and the RBS of the *yabJ* and *spoVG* mRNA are detailed relative to the *yabJ-spoVG* transcription start site (+1). An RNA-RNA base-pair interaction is depicted by an adjoining purple line. (**D**) *In vitro* transcribed RNA fragments used for EMSA analyses corresponding to the yabJ 5’UTR and *yabJ*-*spoVG* RNA. The nucleotide genomic position representative of the *yabJ-spoVG* transcription start site (+1 site) are indicated *below*. The position of each hybrid read recovered from RNase III-CLASH (highlighted in blue and orange), and antisense oligonucleotides (80-mers) used for EMSAs are indicated. (**E**) EMSA analysis of the radiolabelled *yabJ*-*spoVG* intercistronic region (designated as *yabJ*-*spoVG*) and the *yabJ* 5’UTR. (**i**) Increasing concentrations (0, 50x or 100x) of antisense oligomer A or B was titrated with 2.5 pmol *yabJ* 5’UTR against 50 fM of radiolabelled *yabJ*-*spoVG*. Free radiolabelled *yabJ*-*spoVG* is indicated by the black arrow. (**ii**) Quantification of the ratio of free radiolabelled *yabJ*-*spoVG* RNA with and without 2.5 pmol *yabJ* 5’UTR by densitometry.

**Supplementary Figure 2.**
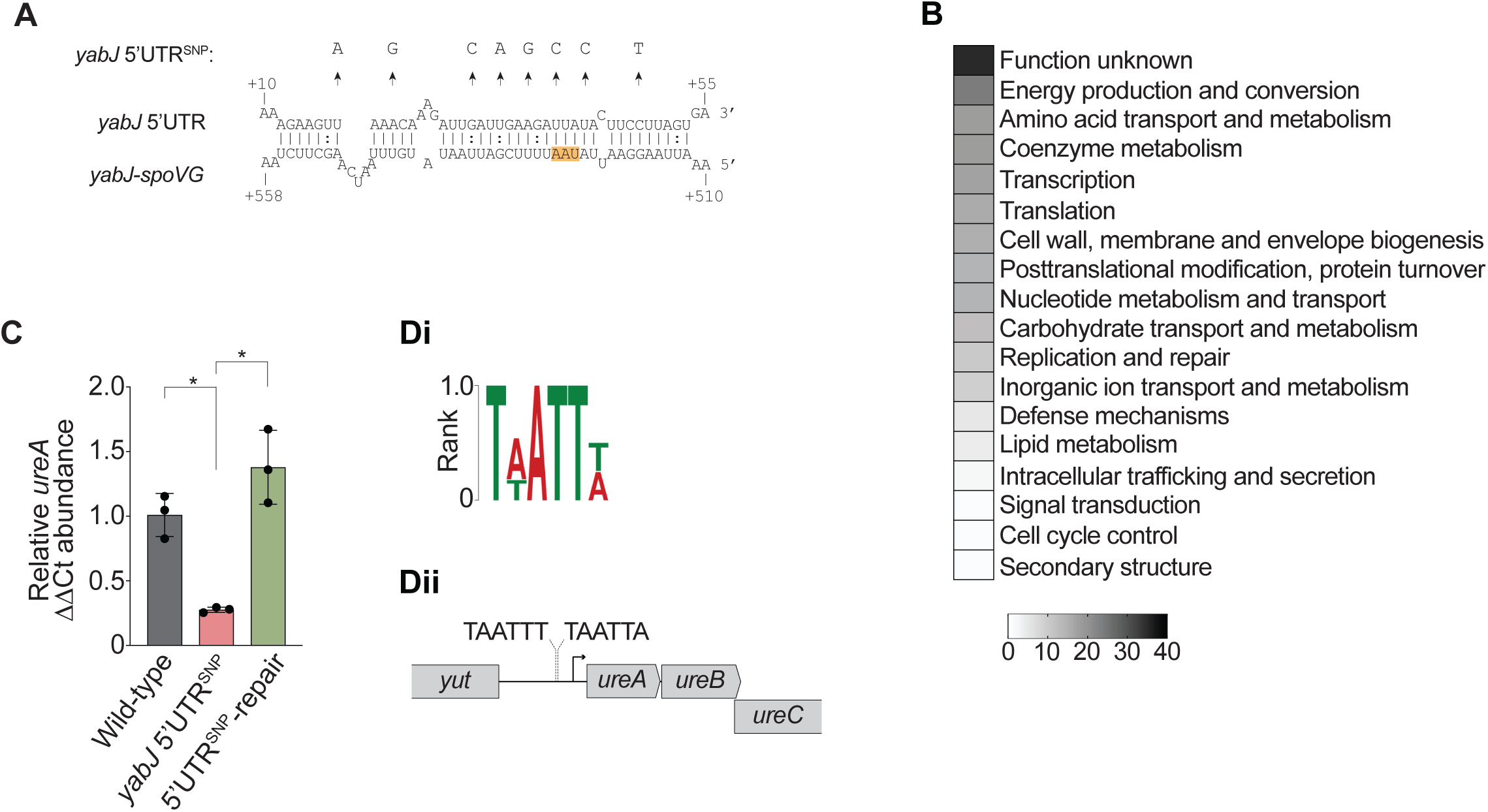
(**A**) Representation of the synonymous point mutations within the *yabJ*-*spoVG* interaction seed used to generate *yabJ* 5’UTR^SNP^ in VISA JKD6008. The nucleotide genomic positions are indicated (*below*) representative of the *yabJ-spoVG* transcription start site (+1 site). (**B**) Clusters of orthologous (COG) classes detailed for differentially abundant proteins from LC-MS/MS analyses (*p*<0.05). The most abundant COG classes are listed in numerical order. (**C**) Histogram of qRT-PCR to quantify *ureA* mRNA chromosomal expression (relative to the *gapA* mRNA) in wild-type, *yabJ* 5’UTR^SNP^, and *yabJ* 5’UTR^SNP^-repair strains. Error bars represent standard error (*n*=3). A student’s *t*-test, two sample assuming unequal variance was used to determine statistical significance. \**p*<0.05. (**D**) Consensus sequence motif for SpoVG DNA-binding identified within MRSA JKD6009 determined using GLAM2SCAN (**i**), and the genomic location of the consensus motifs in the *ureA* transcript (**ii**).

**Supplementary Figure 3.**
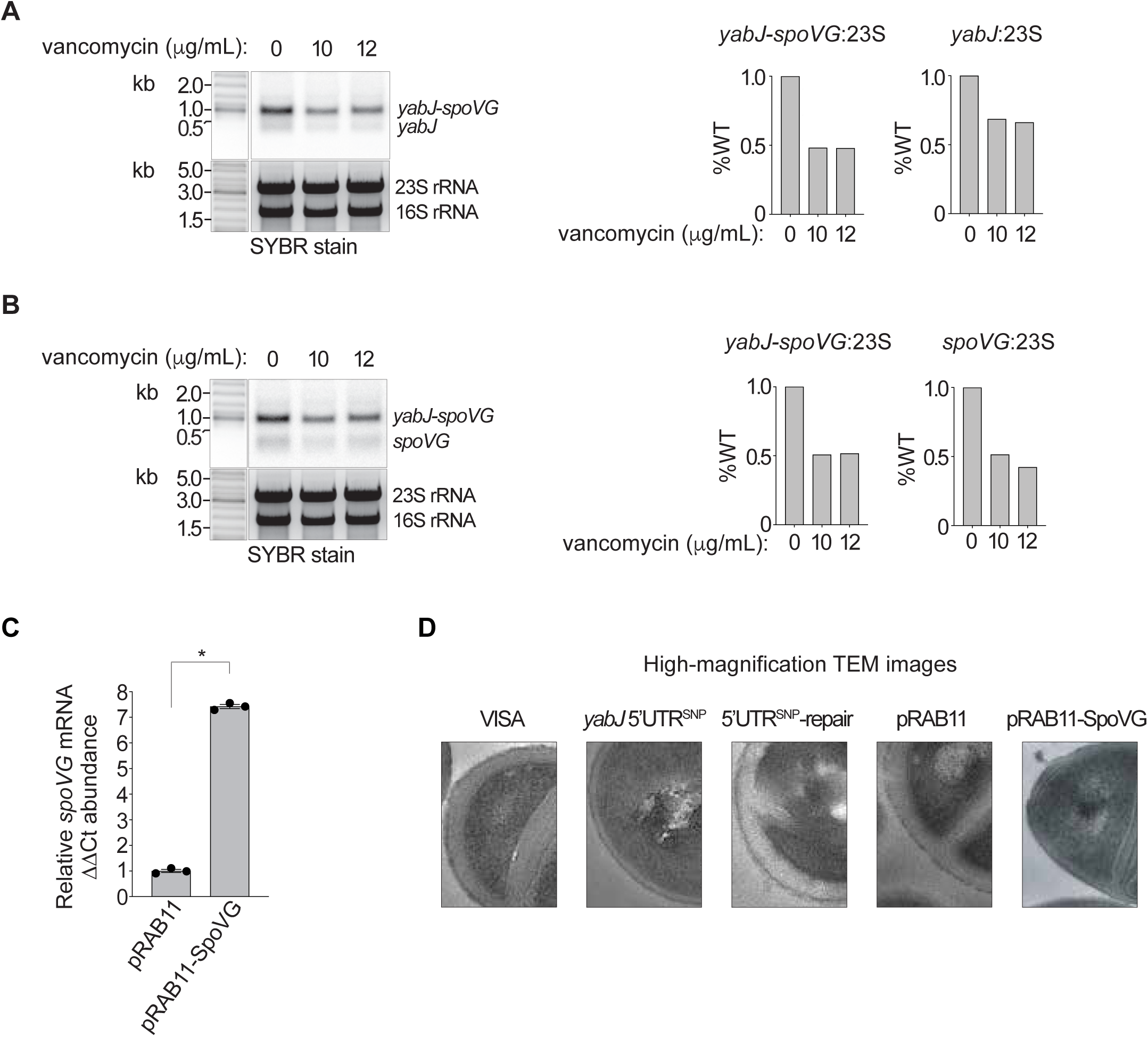
Northern blot analyses of the bicistronic *yabJ*-*spoVG* transcript, and monocistronic *spoVG* and *yabJ* mRNAs. Total RNA was extracted from VISA strain JKD6008 (wild-type) grown to mid-log phase (OD_578nm_ of 2.0) and treated with or without increasing concentrations of vancomycin (0, 10 or 12 μg/mL) for 1 h, and probed for *yabJ* (**A**) or *spoVG* (**B**) transcript. SYBR Green stained rRNA is indicated below as a loading control. Quantification of the ratio of 23S rRNA to either bicistronic *yabJ*-*spoVG*, *spoVG*, or *yabJ* by densitometry, relative to wild-type (*right*). The absence or presence of vancomycin treatment is indicated *below*. (**C**) Histogram of qRT-PCR to quantify the *spoVG* transcript chromosomal expression (relative to the *gapA* mRNA) in VISA JKD6008 pRAB11 and JKD6008 pRAB11-SpoVG strains. Error bars represent standard error (*n*=3). A student’s *t*-test, two sample assuming unequal variance was used to determine statistical significance. \**p*<0.05. (**D**) Close-up images of the cell wall taken using transmission electron microscopy (TEM) of VISA JKD6008 and derivative strains.

